# Effect of Feedback Valence on Family Resemblance Category Learning

**DOI:** 10.1101/764738

**Authors:** Jing-Shang Che, Qiang Xing, Ai-mei Li

**Author notes:** Address correspondence to: Qiang Xing. Department of Psychology, Guangzhou University, NO.230 Waihuan West Road, Guangzhou Higher Education Mega Center, Guangzhou(P.R. China) 510006, (Q. Xing).

## Abstract

This research uses ERPs method to study how different valence of feedback influence family resemblance category learning. The results showed that at the behavioral level, participants in the negative feedback condition received higher test scores than those in the positive feedback condition; at the physiological level, the four kinds of ERPs evoked by both negative and positive feedback are P200, P300, and FRN. Compared with the non-feedback condition, P300 is more sensitive to negative feedback, and negative feedback induces larger amplitude, while P300 is not sensitive to positive feedback. For both negative and positive feedback, when feedback was presented after 200-300ms, the reaction to errors induced FRN production, and FRN produced under negative feedback condition led to more activation. This research has deepened our understanding of the influence of feedback valence on category learning from the brain mechanisms as well.

## Introduction

In real life, we tend to classify objects into different categories based on similarities among objects or rules, and the ability to categorize objects can be learned in different ways (Xing & Sun, 2017). Classification learning as a primary way of category learning has its fundamental paradigms: during the whole process, the participants are required to categorize a complete set of stimuli, which are presented one by one, and with the help of feedback from their responses, the participants are able to obtain the knowledge pertaining to categories (Johansen & Kruschke, 2005). Thus, it can be seen that feedback is essential for classification learning (Sun & Xing, 2014).

According to feedback valence, it can be divided into positive feedback and negative feedback. Positive feedback refers to giving feedback to the correct response, and negative feedback refers to giving feedback to the wrong response (Ashby & O’brien, 2007; Ashby & Maddox, 2010). Previous studies have suggested that the function of feedback is not only the positive reinforcement of the correct response, but also the negative reinforcement of the wrong response. Negative feedback with positive feedback in category learning play a different role, in particular, the positive feedback to the positive reinforcement of correct reaction, but negative feedback focuses on the negative reinforcement of the error response, and the need to synthesis reasoning on the different characteristics of category (Kulhavy & Anderson, 1972; Monchi, Petrides, Doyon, Postuma, Worsley, & Dagher, 2004; Xing, Sun, &Che, 2015). However, Smith and Kimball (2010) believe that negative feedback has a greater role than positive feedback. Because the feedback role is to encourage learners to find the right answer in category learning, making mistakes provide learners with more information, and telling the learners what characteristics is irrelevant. In addition, error response provided by the signal to guide the rules and the choice of strategy transformation. Then, how do positive feedback and negative feedback affect category learning? With the development of cognitive neuroscience technology, compared to traditional behavior experiment, event related potentials (ERPs) is quite sensitive to many cognitive events, which provides a good method to explore effect of positive feedback and negative feedback on category learning.

Cognitive neuroscience research has found that P200, P300, and FRN are involved in feedback processing(Hajeak, Moser, Holroyd, & Simons, 2006; Polezzi et al., 2008). Polezzi et al. found that P200 is sensitive to expected feedback and that unexpected feedback can induce a greater P200 than expected feedback. Within 300~400 ms after feedback is presented, the study found the presence of P300. P300 has been generally considered as the ERP component associated with cognitive functions, such as attention, decision-making, and results assessment (Sato et al., 2005; Yeung & Sanfey, 2004). The amplitude of P300 is proportional to the degree of cognitive resources input; it can reflect how the brain allocates attention resources (Leng & Zhou). Previous research has considered P300 to be sensitive to the feedback number (such as what level of rewarding information, like a reward of + 3 or a reward of + 1, might be given) but free from the influence of feedback valence (i.e., neither positive nor negative feedback) (Polezzi, Lotto, Daum, Sartori, & Rumiati, 2008).

In addition, studying event-related potentials (ERPs) analysis of concurrent feedback stimuli have shown that 200-300 ms after feedback is presented, negative feedback, compared with positive feedback, activates a more drastic negative wave, known as feedback-related negativity (FRN) (Li & Li, 2008). Then, Yeung and Sanfey has shown that FRN is sensitive to result feedback (either positive feedback or negative feedback) but is not sensitive to the amount of reward and punishment; in other words, FRN simply evaluates an event as good or bad. FRN may reflect a simple and quick evaluation of “good” or “bad” with regard to the consistency between results and expectation. Negative feedback (such as responses to error or loss of money) amplifies FRN more than positive feedback does, which shows that the sensitivities of FRN differ in degree to positive and negative feedback.

FRN not only tells whether the results are “good” or “bad” but also provides reflection on how to adjust behavior in the future. Cohen and Ranganath (2007) found that the magnitude of FRN can predict whether or not the participants will adjust their behavior or change strategy next time. Van der Helden et al. (2010) also found that when participants learn from current negative feedback, they will choose an option the next time which was not previously chosen (i.e., successful adjustment behavior) and with more drastic FRN volatility; however, when participants fail to learn from negative feedback, they tend to select the previous option (i.e., they will repeat the wrong option), with less drastic FRN volatility.

However, these conclusions have been derived mostly from gambling tasks. The feedback effect in these studies involves the amount of compensation or reward and punishment mechanisms, and there may be a complex emotional process (Cui & Zhang, 2013). On the other hand, in real life, we carry out classification based mainly on family resemblance, which refers not only to certain similarities in characteristics between or among family members, but also to differences in specific cases, which is an important basis for classification (as showed in Table 1). Category learning based on family resemblance emphasizes the similarities between or among members of the category (Yamauchi & Markman, 1998). Compared to category learning based on rules and information integration, family resemblance material has much in common with natural category materials, and participants can use various strategies for learning (Xing, Sun, & Che,2015). As summarized above, the current studies concerning feedback are based mainly on punishment-reward research, whereas the effects of positive versus negative feedback in pure cognitive feedback in the family resemble category of learning have not been examined.

**Table 1.**
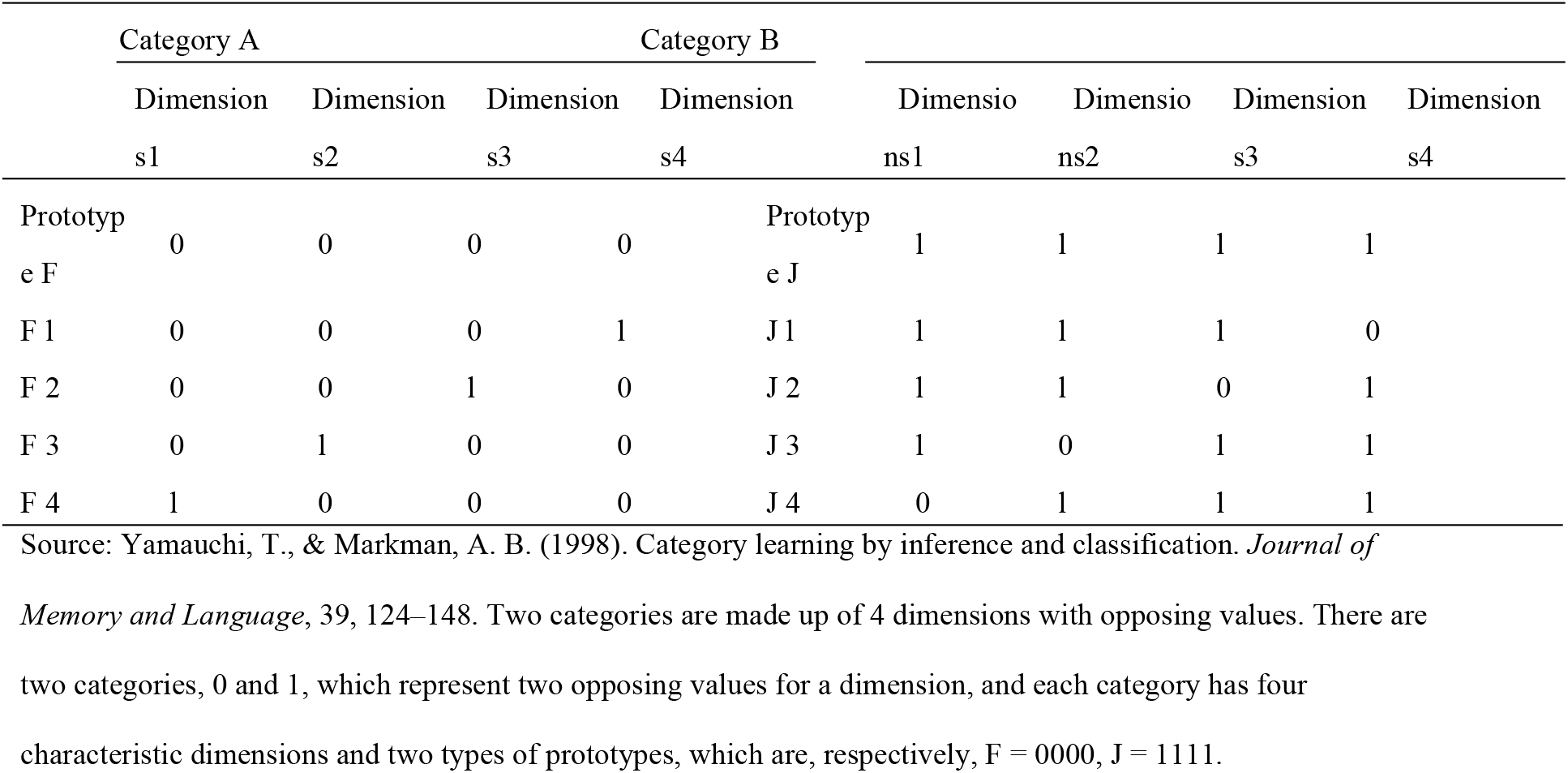
Yamauchi and Markman’s (1998) use of the category structure

Therefore, this study intends to discuss the effect of feedback valence on family resemble category learning and its cognitive neural mechanism by using ERPs. Specifically, according to the feedback valence, there are the positive feedback group and the negative feedback group, the positive feedback condition provides positive feedback when participants have correct response, while the response error does not provide any feedback. Negative feedback condition is similar. Due to different feedback conditions, the mechanism of feedback in category learning is discussed by analyzing specific brain electrical components.

## Method

### Participants

Twenty right-handed college students (10 females and 10 males, M age = 22.97 years, SD = 5.87) were recruited from the Guangzhou University. The participants were divided into two conditions: positive feedback condition and negative feedback condition—and each group was randomly assigned 10 participants. If the accuracy of three consecutive experimental blocks reached 90%, we can say the learning was successful, whereas if the accuracy of thirty blocks still did not reach 90%, then the learning was viewed as unsuccessful. We excluded those participants who failed to manage learning successfully during the experimental process. Thus, we had 13 valid participants (positive feedback condition is seven participants, and other condition is six participants). Data from one participant were discarded due to EEG artifact problems. Participants had no history of head injury and were not taking psychotropic medications. All participants provided informed consent prior to the experiment. Data from the remaining twelve participants were statistically analyzed using an EEG.

### Materials

According to the related dimension presented in Table 1, we used Photoshop to design pictures, setting the pixels to 1024×768. The picture is composed of a category label (F or J) and four dimensions. The prototypes of F and J are shown in Fig. 1.

**Fig. 1.**
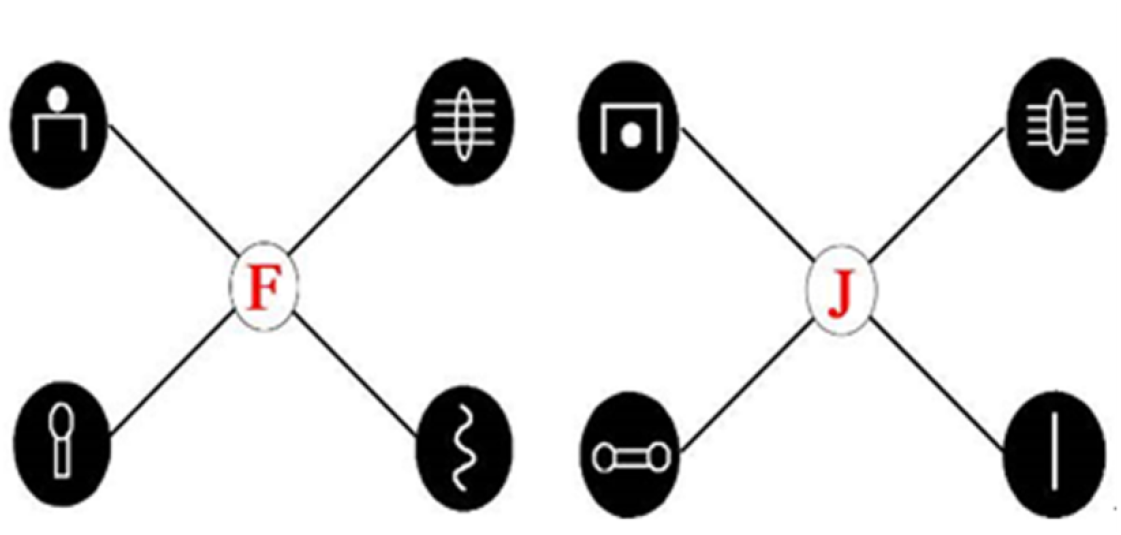
prototype of F and J

### Experimental Design

The experimental design was confined to a single factor between subjects. The independent variable is the feedback valence, and its two levels are positive feedback and negative feedback. The dependent variables include the number of blocks required to meet the standards and the accuracy in the learning and testing phases. In the ERP system, four labels can be made according to whether or not the participants’ responses are provided with feedback. These are S44 (positive feedback, correct), S55 (positive feedback, error), S66 (negative feedback, error), and S77 (negative feedback, correct). In the positive feedback group, if a participant’s correct response was provided with positive feedback, then the instance can be marked S44, whereas S77 can be used if a participant’s erroneous response doesn’t receive any feedback.

### Procedure

E-Prime 2.0 software was adopted for programming, and the display was presented at 1024×768 resolution ratio. The learning-testing paradigm was used in this experiment.

In the learning phase (as showed in Fig. 2), stimuli were presented first; the participants then tried to determine to which category each of these stimuli belong, and, finally, feedback was provided. Prototype stimuli were not presented in the experiment, whereas non-prototype experimental stimuli were randomly presented. Eight trials were counted as one block, and there was an interval of one minute for rest after every three blocks. If a participant’s accuracy was at least 90% in three consecutive block tests, he or she could pass to the testing phase (the learning phase was also ended if a participant’s accuracy could not meet the learning standard in 30 blocks). The two conditions of negative feedback and positive feedback were presented as follows: a red marker, “ √” was shown for a correct judgment under the positive feedback condition; otherwise, nothing was shown. A red "× " appeared for an incorrect judgment in the negative feedback condition; otherwise, nothing was presented.

**Fig. 2.**
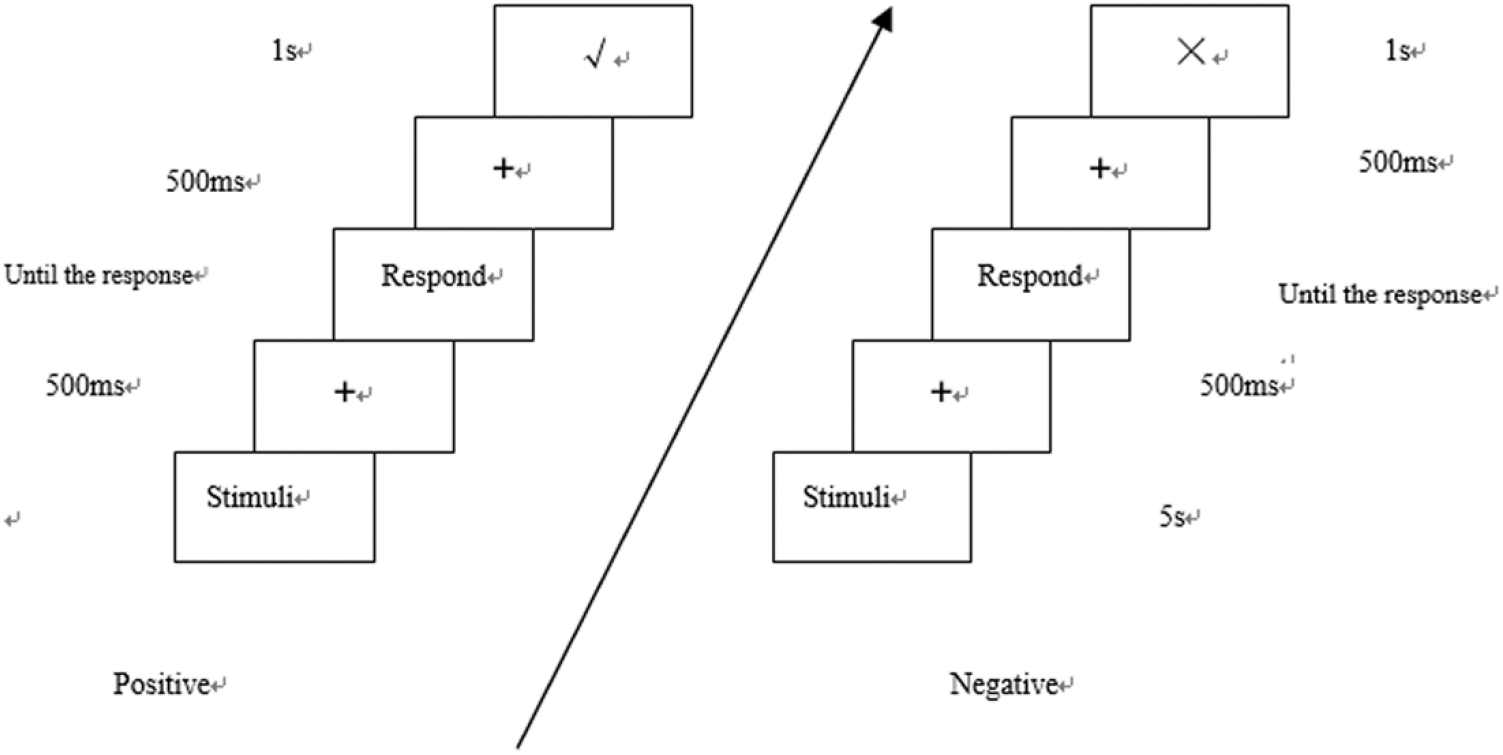
Positive feedback vs. negative feedback experiment flow chart

Following the experiment design of Liu and Mo (2008), the testing phase consisted of three sub-tests, in which feedback was not provided. The first one was the classification test, wherein participants were asked to categorize pictures presented as in a learning phase, among which two categories of prototype had been added. The second one was the memory test (Fig. 3a), which asked participants to determine to which category every single characteristic belongs. The third one was a reasoning test (Fig. 3b), in which the participants were asked to determine which characteristic presented on the dotted line belongs to the same category as the picture. Achievement in these three test modes is affected by the depth of learning, and classification learning is primarily about simple memory because its content is presented in the learning stage. The memory test is about remembering every single feature, whereas for the reasoning test, in order to have a better performance, participants needed to reason about internal relationships inside the category or category structure. Using three different tests, though they differed in the depth of processing, helped us to study thoroughly the characteristics of positive feedback and negative feedback.

**Fig. 3.**
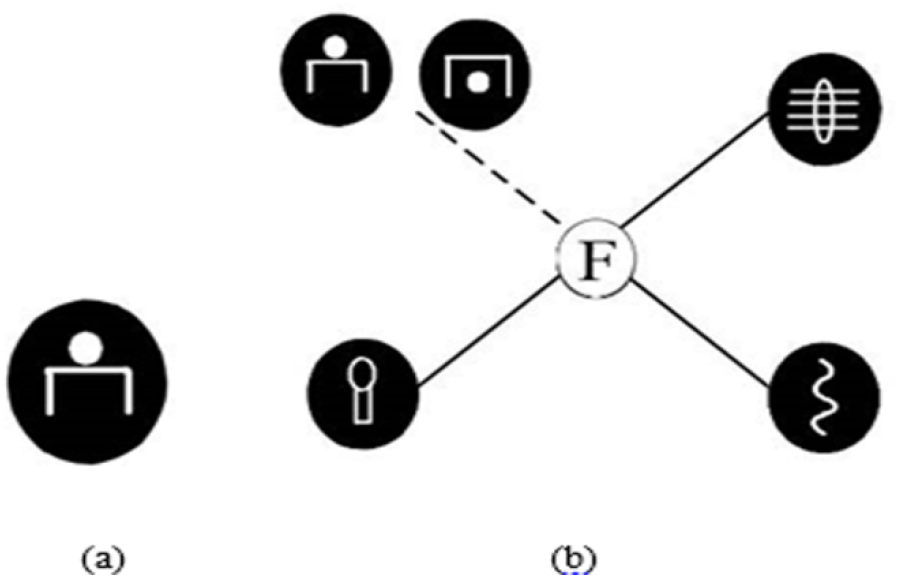
Single-feature testing (a) and reasoning test (b) sample

### Data recording and Analyzing

The EEG was recorded from an array of 32 electrode sites placed in accordance with the extended International 10–20 System using an elastic cap (Brain Product, GmbH, Germany). Voltages were amplified by low-noise electrode differential amplifiers with a frequency response of DC 0.01–80 Hz and digitized at a rate of 500 samples per second. The digitized signals were recorded with Brain Vision Recorder software. The horizontal electrooculogram (HEOG) was recorded with two electrodes placed laterally to the right and left eyes. The vertical electrooculogram (VEOG) was recorded with electrodes placed above and below the right eye. All inter-electrode impedances were maintained below 5 KΩ. All EEG signals were with an online reference to the left mastoid. Trials containing EEG sweeps with amplitudes exceeding ±80 μV were excluded.

We adopted Analyzer 2.0 software to analyze the data. For each stimulus, epochs of 1000 ms in duration, including a 200 ms pre-stimulus period used as baseline, were extracted from the continuous EEG record. In using the stimulus type of markers, the segmentation function is used to do segmentation, taking feedback incentives as the zero point for superimposition to obtain a total average figure (Zhao, 2004). Repeated measures analyses of variance (ANOVAs) for all ERP components were conducted for feedback valence (negative emotion vs. positive emotion) by feedback (have or not) as between-participants factors. A within-participants factor was the electrode. Based on previous literature and visual inspection of the ERP grand average waveforms for all conditions (Polezzi et al., 2008; as showed in fig.4 and fig.5) (Kok, 2001; Polich, 2007; Folstein and Van Petten, 2008). According to the experimental purpose and the overall average figure, six electrode points were selected to be analyzed, which are Fz, Fc1, Fc2, Cz, Cp1, and Cp2. Frontal P200 component was measured as a peak amplitude within the post-stimulus time windows of 100 to 220 ms, and P300 component was measured as the mean amplitude during 300 to 400 ms, and FRN (peak amplitude during 200–300 ms) components were measured on electrodes Fz, Fc1, Fc2, Cz, Cp1, and Cp2.The resulting p values are corrected by using the Greenhouse-Geisser.

## Results and Analysis

### Behavior results

In the learning phase, participants in the negative feedback condition needed (13.83 ± 7.99) blocks to reach the learning standard, but for participants in the positive feedback condition, (19.43 ± 8.79) blocks were needed. The learning speed of participants in the positive feedback condition was slower than that of participants in the negative feedback condition, but an independent samples T-test showed no significant difference between them, t (11) = −1.192, p > 0.05.

In the test phase, concerning the classification testing performance, the accuracy of participants who received positive feedback is (0.94 ± 0.23), and that of participants who were given negative feedback is (0.98 ± 0.13); there is no significant difference between them, t (11) = 1.194, p > 0.05. In the memory testing performance, the accuracy of those who received positive feedback is (0.82 ± 0.39), and that of participants who were given negative feedback is (0.90 ± 0.31); there is also no significant difference between those two figures, t (11) = 1.072, p > 0.05. As for the reasoning test, the accuracy of participants who received positive feedback is (0.90 ± 0.31); that of participants who were given negative feedback is (0.74 ± 0.44). The difference between them is huge, t (11) = 4.205, p < 0.001, indicating that the participants in the negative feedback group had better performance in reasoning than those in the positive feedback group.

### ERP results

In order to test whether there is a difference among EEG components induced during the process of categorization after different feedback conditions were presented according to four types—namely, “positive feedback condition-correct response”, “positive feedback condition-wrong reaction”, “negative feedback condition-correct response”, and “negative condition - wrong reaction”—this study classified and superimposed the EEG to get the ERPS of four different conditions. Under “positive feedback condition-correct response” conditions, the average effective overlay time is 79; under “positive feedback condition-wrong reaction” conditions, 42; under “negative feedback condition-correct response” conditions, 57; under “negative feedback condition-wrong reaction” conditions, 40. At the same time, the total average figure shows that P200, P300, N500, FRN, and other EEG components evoked have sharp peaks, so using the method of maximum amplitude, we can further statistically test each component through an analysis of amplitude and latency.

### P200

Under conditions wherein an error response was provided with both positive and negative feedback groups, P200 was activated, which can be seen in Fig. 4. For the latency of P200, within the time window of 100~220 ms, we conducted 2 (negative feedback condition-wrong reaction, positive feedback condition – wrong reaction) × 6 electrodes (Fc1, Fc2, Cp1, Cp2, CZ, Fz) repeated ANOVAs, with no main effect and interaction effects appearing. This indicates that both positive and negative feedback induced P200, but no significant differences were shown concerning when the inducement happened.

**Fig. 4.**
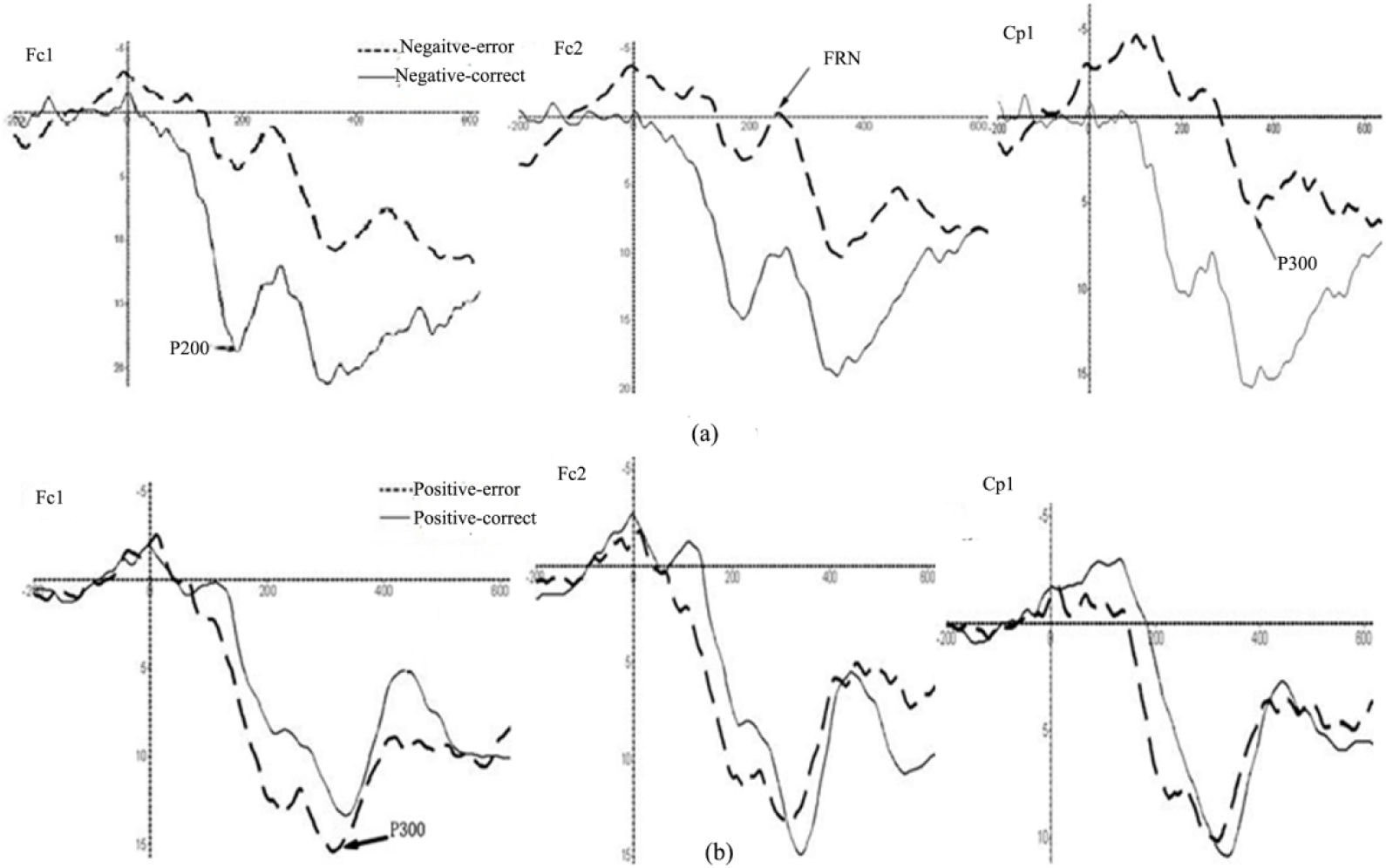
(a) the total average figure for negative feedback condition-error and positive feedback condition-error. (b) The total average figure for positive feedback condition-correct and negative feedback condition–correct.

For the peak value of P200, we conducted 2 (negative feedback condition-wrong reaction, positive feedback condition – wrong reaction) × 6 electrodes (Fc1, Fc2, Cp1, Cp2, Cz, Fz) repeated ANOVAs. The result reveals that the main effect of feedback is significant, F(1,10) = 9.13, p = 0.013, η2 = 0.78, and the amplitude appearing under a negative feedback condition is significantly less drastic than that under conditions in which no feedback was provided; the main effect of the electrode is significant, F(5,50) = 13.01, p = 0.001, η 2 = 0.97, but interactions between feedback and electrodes are not significant, F(5,50) = 1.72, p = 0.21.

### P300

#### (1) Participants’ error responses

Under conditions in which an erroneous response was provided with positive feedback and negative feedback, within the time window 300~400 ms and for the latency of P300, we conducted 2 (negative feedback condition-wrong reaction, positive feedback condition – wrong reaction) ×6 electrodes (Fc1, Fc2, Cp1, Cp2, Cz, Fz) repeated ANOVAs; the result shows that the main effects and interactions are not significant.

For the peak value of P300, we conducted 2 (negative feedback condition-wrong reaction, positive feedback condition – wrong reaction) × 6 electrodes (Fc1, Fc2, Cp1, Cp2, Cz, Fz) repeated ANOVAs. The result shows that the main effect of the feedback is significant, F(1,11) = 11.83, p = 0.006, η2 = 0.87; the main effect of the electrode is not significant, F(5,50) = 3.13, p = 0.084; and interactions between feedback and the electrode are not significant, F<1.

#### (2) Participants’ correct responses

Under conditions in which a correct response was provided with both positive and negative feedback within the time window of 300~400 ms and for the latency of P300, we conducted 2(negative feedback condition-right reaction, positive feedback condition – right reaction) ×6 electrodes (Fc1, Fc2, Cp1, Cp2, Cz, Fz) repeated ANOVAs; the result shows that the main effects and interactions were not significant.

For the peak value of P300, we conducted 2 (negative feedback condition-right reaction, positive feedback condition – right reaction) ×6 electrodes (Fc1, Fc2, Cp1, Cp2, Cz, Fz) repeated ANOVAs. The result shows that the main effect of feedback is not significant, F<1; the main effect of the electrode is significant, F(5,50) = 8.79, p < 0.001, η2 = 0.98; and the interactions between feedback and electrodes are not significant, F(5,50) = 1.04, p > 0.05.

#### (3) Positive feedback and negative feedback

Under conditions in which a correct response was provided with positive feedback, as well as those under which an error response was provided with negative feedback within the time window of 300~400 ms and for the latency of P300, we conducted 2 (negative feedback condition-wrong reaction, positive feedback condition – right reaction) ×6 electrodes (Fc1, Fc2, Cp1, Cp2, Cz, Fz) repeated ANOVAs; the result shows that the main effects and interactions are not significant.

For the peak value of P300, we conducted 2 (negative feedback condition-wrong reaction, positive feedback condition – right reaction) ×6 electrodes (Fc1,Fc2, Cp1, Cp2, Cz, Fz) repeated ANOVAs; the result shows that the main effect of feedback is not significant, F(1,10) = 1.70, p > 0.05; the main effect of the electrode is significant, F(5,50) = 5.44, p = 0.008, η2 = 0.84; and interactions between feedback and the electrodes are not significant, F(5,50) = 1.36, p > 0.05.

### FRN

Under the error response condition within the time window of 200 ms~300 ms and for the latency of FNR, we conducted 2 (negative feedback condition-wrong reaction, positive feedback condition – wrong reaction) × 6 electrodes (Fc1, Fc2, Cp1, Cp2, CZ, Fz) repeated ANOVAs; the result showed that the main effects and interaction effects are not significantly different, regardless of whether positive or negative feedback is given to an error response in the incubation period.

For the peak value of FRN, we conducted 2 (negative feedback condition-wrong reaction, positive feedback condition – wrong reaction) × 6 electrode (Fc1, Fc2, Cp1, Cp2, Cz, Fz) repeated measures ANOVAs. The result shows that the main effect of feedback is significant, F(1,10) = 8.02, p = 0.018, η2 = 0.72; the main effect of an electrode is not significant, F(5,50) = 2.20, p = 0.157; and interactions between feedback and electrodes are not significant, F<1.

## General discussion

Taking family resemblance category as the object of study, this research investigated the electrophysiological mechanisms that relate to how feedback affects category learning. The behavioral results show that, compared with the effect of positive feedback, negative feedback plays a more important role. First, with regard to learning speed, though the differences are not very significant, statistically speaking, the participants’ learning speed in the negative feedback group trended toward being faster than that in the positive feedback group (19.43 > 13.83). Second, comparing the performance in the test phase under the positive feedback condition with that under the negative feedback condition, the results showed that for the classification and memory tests, the participants in both conditions performed well with regard to the judgment they had done in the learning phase. In the reasoning test, which required the participants to learn in depth about category structure in order to achieve a better performance, the participants in the negative feedback condition showed a significantly better performance than those in the positive feedback group, indicating that it is negative feedback that may help participants the most in further processing the stimulus and during which the inference rule strategy may be employed.

The EEG data shows that within 100~220 ms after feedback was presented, both positive and negative feedback induced the production of P200; within 200~300 ms after the presentation of feedback, the correction of P300 is more likely to be induced under negative feedback than when no feedback is presented, but there is little difference between the amplitudes induced by both positive and negative feedback. When either positive or negative feedback is given to error responses, FRN can be induced within 200~300 ms after presentation of the feedback, and the volatility of FRN under negative feedback is much more drastic than that under positive feedback.

### P200

Polezzi et al. (2008). indicates that P200 is sensitive to expected feedback but that unexpected feedback can induce a bigger P200 than expected feedback. Our experiment found that if no feedback is provided, P200 can be better activated than if negative feedback is given. The former condition, by contrast with negative feedback, is much closer to unexpected feedback.

Because this study did not use the probability feedback, the positive feedback group would respond to the correct response of the participants, and the wrong response would be blank screen, and the negative feedback group would be similar. Since each stimulus has only errors and the correct two situations, it may be possible to guess whether the response is correct or not in the absence of feedback. However, under different conditions, the induced P200 components can be found that the amplitude of P200 is significantly greater than that in the non-feedback condition (i.e. blank screen). If being try to guess the show blank screen for response error, so blank screen appears is the negative feedback, but from the eeg data can be found on the blank screen conditions (i.e., no negative feedback) and there is negative feedback between the group of P200 amplitude difference reached significant level, to some extent, illustrates the subjects did not guess blank screen presents its own response correctly or not.

Considering the research purpose, we did not use probability feedback for this study: namely, that the positive feedback group would receive feedback for each correct response, and the case was similar for the negative feedback condition. Here, a problem may arise: because each stimulus is either correct or erroneous, even if no feedback is provided, participants are able to infer whether their responses are correct or in error. However, if participants are able to infer that the responses are correct or erroneous without receiving any feedback, then the waveform of the positive feedback condition is similar to that of the negative group, but the waveforms in Fig. 4 and 5 showed that significant differences among waveforms appeared under conditions in which neither positive nor negative feedback was provided, indicating to some extent that participants’ guessing is less likely to influence the results.

**Fig. 5.**
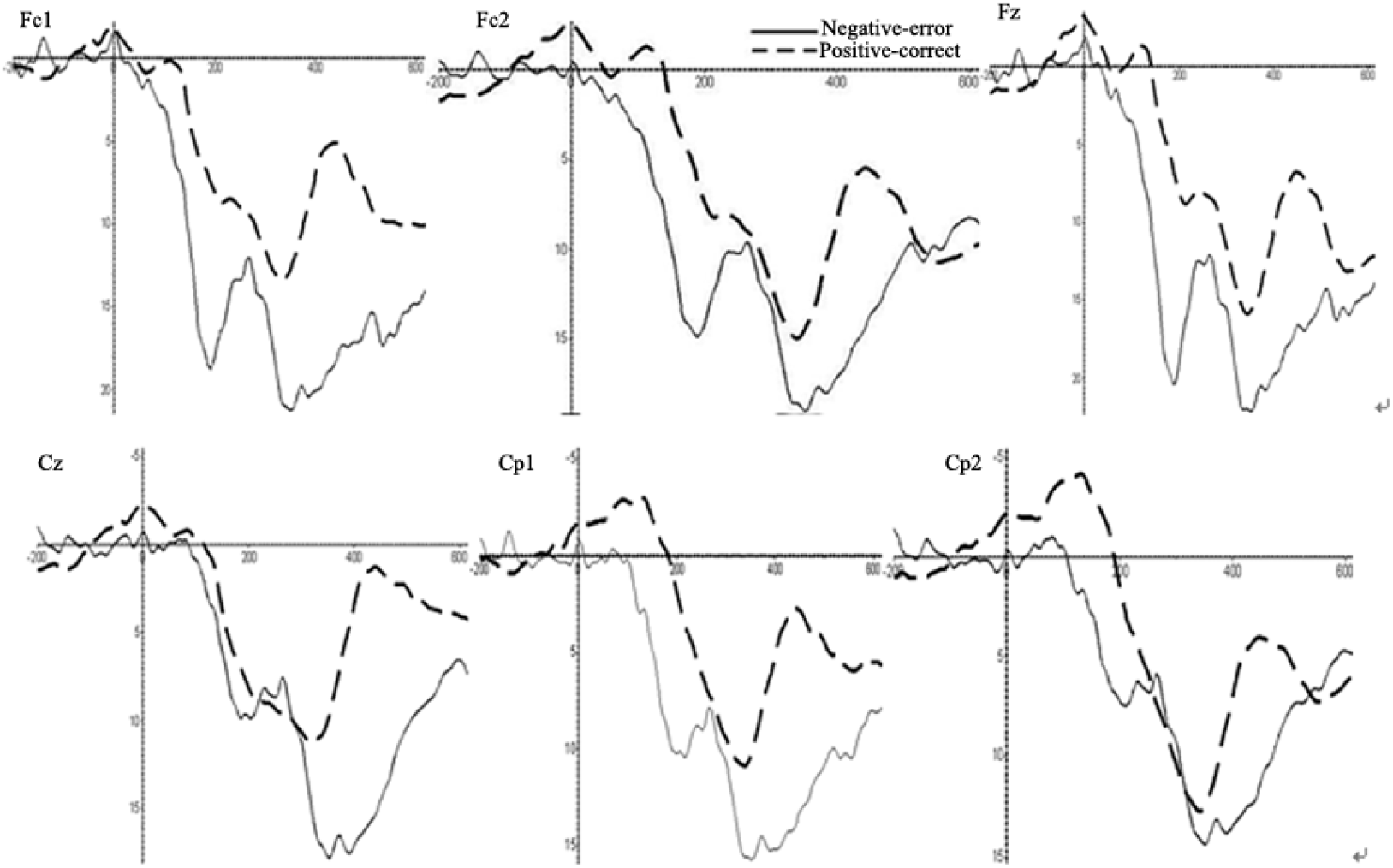
The total average figures under positive feedback condition-correct and negative feedback condition-error.

### P300

Previous studies have shown that P300 has something to do with memory updates, encoding, and retrieval, and the amplitude fluctuation of P300 reflects the degree of input of attention resources (Donchin & Coles, 1988).

In this study, P300s were activated under both negative and positive feedback, which fully explains that the memory system plays an important role in category learning based on feedback. Compared with no feedback, P300 is more sensitive to negative feedback. This condition induced a more drastic volatility, indicating that learners can be more easily attracted to devoting more attention resources to further processing if negative feedback, rather than none at all, is provided. At the same time, we found that for the family resemblance category structure, both positive and negative feedback activated P300, but there was no significant difference in the degree of activation. A study by Yeung and Sanfey(2004) has shown that P300 may be related to the degree to which emotions are awakened. Liu, Tang and Ye (2012) suggest that P300 can induce a more drastic volatility under rewarding conditions. For pure cognitive feedback, the emotional awakening from either negative or positive feedback plays a less important role than a rewarding task does.

### FRN

Within 200~300 ms after feedback for an erroneous response is provided with either negative or positive feedback, FRN can be induced, but the amplitude under negative feedback was significantly higher than for positive feedback. Van der Helden et al. (2010) also found that when participants learn something from current negative feedback, they will subsequently choose an option they’ve never tried before (i.e., successfully regulate their behavior), resulting in a more drastic amplitude of FRN. Conversely, when participants learn nothing from the current negative feedback, they will choose options that have been selected before (i.e., they repeat wrong options), and in such a situation the amplitude of FRN is less drastic. This indicates that compared with positive feedback, negative feedback can better help participants adjust learning strategies and consequently learn category knowledge better.

However, when a correct response was provided with both positive and negative feedback, we did not find FRN. Holroyd and Coles (2002) argues that when the signal of negative reinforcement learning is transferred to ACC through the midbrain dopamine system, FRN will show itself. If the current event caused by an error response is worse than expected, then the EEG negativity will be amplified more drastically. Given pure cognitive experiments when only right and wrong feedback are provided, if participants think that a response is correct but the feedback informs them that they are wrong, the current event is not better than what they expected. If the situation is reversed, then the current event is better than expected. However, if no feedback is given, participants have no way to know whether their response is correct or erroneous. FRN was found under negative feedback if the response was in error, but it will not appear if the answer is correct under positive feedback. A possible explanation for this is that participants usually expect to be correct, and because positive feedback is consistent with what participants expect, FRN does not show itself. We can also provide a more effective explanation for negative feedback, which induces FRN: negative feedback, which is inconsistent with expectation, inspires participants’ motivation, causing them to pay more attention to stimulus information; accordingly, the negative feedback lends itself to further processing.

### Reinforcement learning or error-driven

According to the reinforcement learning model, positive reinforcement encourages desired behavior, whereas negative reinforcement can reduce the probability of unwanted behavior (Niv,2009). Positive feedback, therefore, can promote the coupling between label and stimulus. This study suggests that compared with positive feedback, negative feedback plays a more important role in learning. Error-driven learning acts as a fundamental driving force that encourages learners to find the inherent rules among objects, in turn helping learners more easily apprehend the essence of objects. Compared to positive feedback, negative feedback can provide more information because the participant knows both what is correct and what is error (Smith, & Kimball, 2010). In this study, under positive-correct and negative-correct conditions, the amplitude value of P300 is larger under negative feedback, indicating that it can attract learners to invest more attention resources to enable further processing. For category learning, negative feedback is more effective than positive feedback. On the other hand, under positive-error or negative-error conditions, FRNs were induced. In addition, the amplitude of FRN under negative feedback is significantly more drastic than that under positive feedback, which indicates that individuals have more activation in the cerebral cortex under negative feedback, which contributes to further processing and to adjustment of strategies. From these results, we believe that the negative feedback has a greater effect on category learning than positive feedback, in support of the error-driven mechanism.

In this study, the number of participants meeting the learning standard in the negative feedback group was lower than for the positive feedback group. It may be that negative feedback itself does not necessarily enable participants to learn about categories, whereas positive feedback can guarantee the success of category learning because giving feedback only for correct responses is more likely to be a memory task, and reinforcement plays a relatively greater role in memory. As a result, as long as a participant has enough time, he or she will eventually keep all of samples in mind. As for negative feedback, the inference rule enables participants to process the stimuli further to find the correct rules. In other words, trial-and-error learning does not necessarily enable participants to acquire knowledge, but once the right strategies are adopted, the participants are able to learn quickly, whereas simple memory can enable learning but is less efficient.

## Conclusions

1. compared with positive feedback, negative feedback is significantly better than positive feedback in reasoning test.
2. P200, P300, FRN, N500 are related to the family resemblance category learning; under positive feedback and negative feedback conditions, the degree of activation of P300 is not different, indicating that attention allocation and the use of memory are equally important under positive feedback and negative feedback conditions;
3. under negative feedback condition, there is a greater activation of FRN, which suggests that negative feedback plays a greater role in regulating participants’ subsequent behavior. The result shows that negative feedback is more effective, and error-driven plays a greater role in learning family resemblance material.

